# Generative neural networks separate common and specific transcriptional responses

**DOI:** 10.1101/2021.05.24.445440

**Authors:** Alexandra J. Lee, Dallas L. Mould, Jake Crawford, Dongbo Hu, Rani K. Powers, Georgia Doing, James C. Costello, Deborah A. Hogan, Casey S. Greene

**Affiliations:** Genomics and Computational Biology Graduate Program, University of Pennsylvania, Philadelphia, PA, USA; Department of Systems Pharmacology and Translational Therapeutics, University of Pennsylvania, Philadelphia, PA, USA; Department of Microbiology and Immunology, Geisel School of Medicine, Dartmouth, Hanover, NH, USA; Wyss Institute for Biologically Inspired Engineering, Harvard University, Boston, MA, USA; Department of Pharmacology, University of Colorado School of Medicine, Denver, CO, USA; Center for Health AI, University of Colorado School of Medicine, Denver, CO, USA; Department of Biochemistry and Molecular Genetics, University of Colorado School of Medicine, Denver, CO, USA

## Abstract

Genome-wide transcriptome profiling identifies genes that are prone to differential expression across contexts (“common DEGs”), as well as genes with changes specific to the experimental manipulation. Distinguishing common DEGs from those that are specifically changed in a context of interest allows more efficient prediction of which genes are specific to a given biological process under scrutiny. Currently, commonly differentially expressed genes or pathways can only be identified through the laborious manual curation of highly controlled experiments, an inordinately time-consuming and impractical endeavor. Here we pioneer an approach for identifying common patterns using generative neural networks. This approach produces a background set of transcriptomic experiments from which a null distribution of gene and pathway changes can be generated. By comparing the set of differentially expressed genes found in a target experiment against the generated background set, common results can be easily separated from specific ones. This “Specific cOntext Pattern Highlighting In Expression data” (SOPHIE) approach is broadly applicable to new platforms or any species with a large collection of gene expression data. We apply SOPHIE to diverse datasets including those from human, human cancer, and the bacterial pathogen *Pseudomonas aeruginosa*. SOPHIE identifies common DEGs in concordance with previously described, manually and systematically determined common DEGs. Further, molecular validation indicates that SOPHIE detects highly specific, but low magnitude, biologically relevant, transcriptional changes. SOPHIE’s measure of specificity can complement log fold change values generated from traditional differential expression analyses. For example, by filtering the set of differentially expressed genes, one can identify those genes that are specifically relevant to the experimental condition of interest. Consequently, these results can inform future research directions.

## Introduction

Genome-wide transcriptomics analysis allows investigators to examine how global gene expression changes under the tested experimental stimulus or across different states, individuals or genotypes. When interpreting the results of these analyses, attention tends to focus on controlling false discoveries^1–4^ – i.e. differential gene expression patterns that arise due to noise or variation during measurement. In addition to false discoveries, however, certain genes tend to be commonly differentially expressed across a diverse panel of environmental stresses.^5^ The response of this collection of genes was termed the environmental stress response (ESR). Despite the ESR being described more than two decades ago^5^, compared to false discoveries, less attention has been paid to controlling for these commonly differentially expressed genes (common DEGs). These findings include differential expression changes that are observed across experiments regardless of the experimental manipulation. Both gene-based^5, 6^ and pathway-based^7^ analyses can return common results.

While these common findings are not false discoveries, they provide little contextual information or insight into the biological process being queried as they are observed in many unrelated experiments. Not knowing which discoveries are common versus specific can lead to misinterpretations or lack of specificity in interpreting results, so it is important to account for these different types of findings in addition to correcting for false discoveries.

Controlling for common findings is inordinately time-consuming and therefore limits the use of protocols that would identify them. Current methods rely on manual curation of a background set of experiments to select experiments with consistent experimental design and platform, as well as to use metadata to group samples for downstream statistical analysis. Re-curation is required to derive an appropriate background distribution in a new context, such as when switching to a new measurement platform, applying a different experimental design or analytical approach, incorporating new data, or examining a different organism. These background experiments are analyzed to identify genes and pathways that are common based on the frequency at which they are differentially expressed in the background experiments.^1, 2^ Even when data are readily available, curating and analyzing hundreds of experiments requires a significant time investment to define a compendium of experiments to use as a background.

We introduce an general approach, termed Specific cOntext Pattern Highlighting In Expression data (SOPHIE), that distinguishes between common versus specific transcriptional signals in a selected template experiment using a generative neural network^8^ to simulate a set of background transcriptome experiments. Using a generative neural network allows SOPHIE to automate the analysis of common DEGs. This approach requires enough gene expression data to generate synthetic measurements; however, the data do not need to be curated by experimental design, which removes a usually time-consuming step. Such data are readily available through NCBI Gene Expression Omnibus (GEO)^9^, Short Read Archive (SRA)^10^, European Nucleotide Archive (ENA)^11^, and other repositories. Many datasets are already processed for reuse through projects such as recount2^12^ or ARCHS4^13^. Because SOPHIE relies on generating synthetic data that match a user-selected template experiment, it can be applied to arbitrary downstream analytical workflows, which could be differential expression (DE) analysis, pathway analysis, or other methods, to provide a background distribution of common findings. Furthermore, by using a single template experiment, we only need to define sample groupings for this one experiment as opposed to manually annotating groups for hundreds of experiments. Overall, without the need for manual curation to define a compendium and group samples, SOPHIE can expand lists of genes for follow-up by identifying genes that are context-specific but have subtle signals and are thus understudied in that context. SOPHIE can also filter lists of genes for functional validation by limiting a list of genes to those that are both differentially expressed and highly specific. Overall, SOPHIE’s specificity score can be a complementary indicator of activity compared to the traditional log fold change measure and can help drive future analyses.

We use SOPHIE to identify common DEGs in a human microarray dataset, and the results are consistent with the prior manually curated report using the same human microarray dataset. Next, we find consistent common DEGs using a different human microarray dataset, a cancer cell line dataset, demonstrating that common DEGs are shared across contexts. Furthermore, we also find consistent common DEGs using human RNA-seq data, demonstrating that common differentially expressed genes are shared across platforms too. SOPHIE is also generalizable across organisms as shown by application to the opportunistic bacterial pathogen and model organism, *Pseudomonas aeruginosa* (*P. aeruginosa*). The metabolic choices of *P. aeruginosa* can impact its pathogenicity and using SOPHIE to analyze alternative carbon utilization in *P. aeruginosa*^14^ reveals gene expression changes that are specific to different regulatory levels in the hierarchy of the carbon catabolite repression cascade. This analysis reveals context-specific regulation of arginine metabolism, whose genes would be undetected in a traditional differential expression analysis due to their low magnitude. Based on our SOPHIE results, we hypothesize that these arginine related gene expression changes are specific to some but not all gene perturbations in the carbon catabolite repression pathway that controls alternative carbon utilization. Experimental data support the prediction that arginine catabolism is specifically perturbed by some, but not all mutations of genes, in the pathway. This demonstrates that SOPHIE can successfully identify candidate genes that are specifically relevant to the context of interest, and difficult to uncover through previously developed analysis tools.

## Results

### SOPHIE distinguishes common and specific transcriptional patterns

The main steps for SOPHIE are illustrated in Figure 1A. The first step is to generate a background set of transcriptome experiments, for which we applied ponyo^8^. Ponyo uses a generative neural network, in this proof-of-concept a variational autoencoder (VAE), to generate new samples that match a selected template experiment’s design (in our case the experiment is comprised of a control and one experimental group) by encoding and shifting samples in the latent space while preserving their relative positioning. Intuitively, this latent space translation is akin to simulating an experiment with the same experimental design but studying a different biological process or a different set of conditions. SOPHIE uses ponyo to simulate realistic-looking transcriptome experiments that serve as a background set for distinguishing common versus specific transcriptional signals.

**Figure 1:**
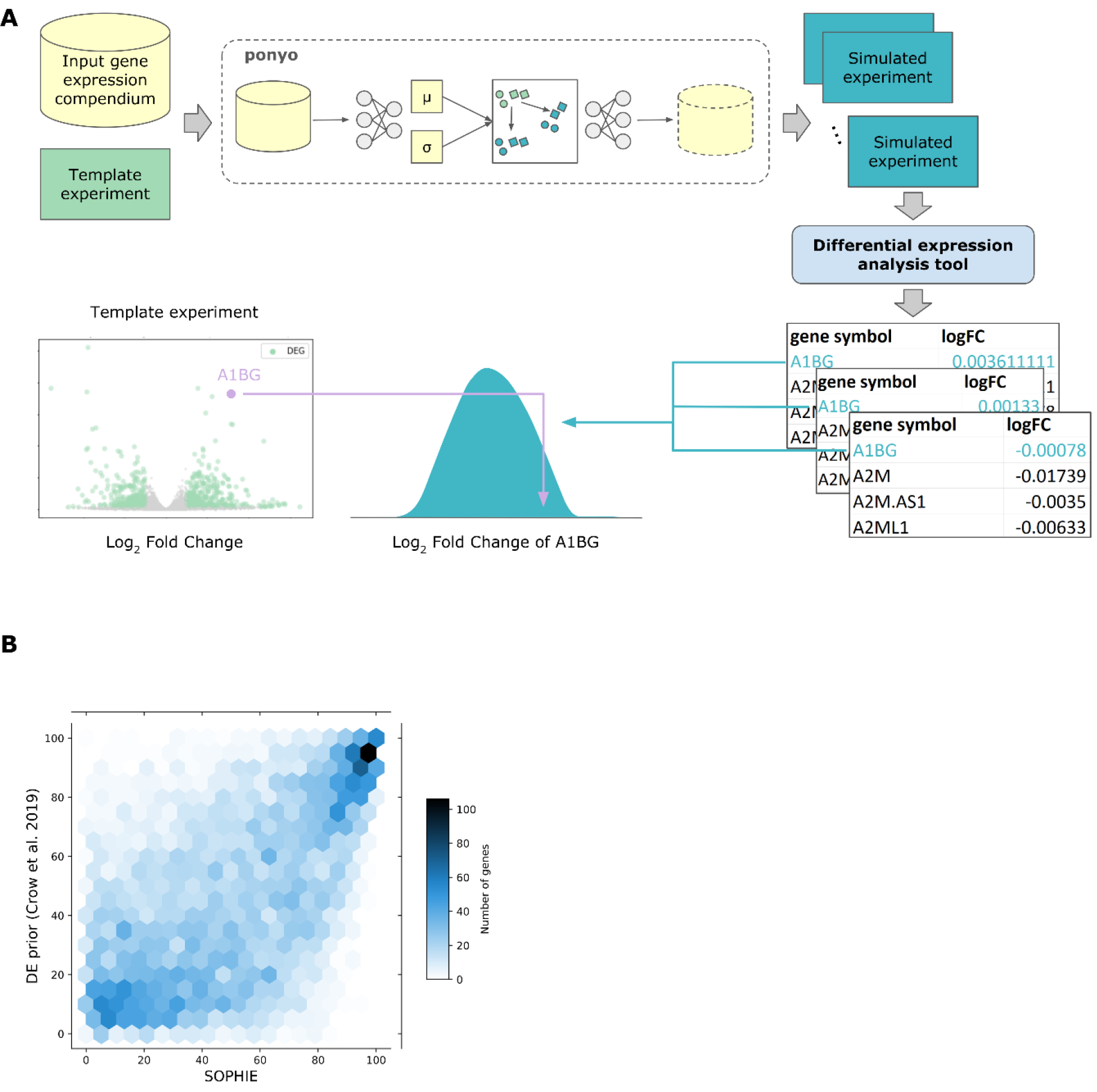
SOPHIE is an approach to distinguish between common and specific DEGs using a generative neural network. A) SOPHIE workflow is designed to distinguish between common and specific transcriptional signals. SOPHIE starts by applying ponyo to simulate gene expression experiments. Next, SOPHIE applies differential expression tools like DESeq2 for RNA-seq data or Limma for array data to get association statistics for each simulated experiment. Finally, SOPHIE returns a distribution of how changed each gene is across the collection of background simulated experiments so that users can compare gene expression changes from their template experiment of interest. B) Spearman correlation between gene percentiles using our SOPHIE approach trained on Crow *et al.* (array) using GSE10281 as a template (x-axis) versus percentiles using manually curated experiments from the same Crow *et al.* (y-axis) had correlation coefficient of 0.59.

For the next step, SOPHIE applies a differential expression analysis tool, like DESeq or Limma, to get association statistics. Then those differential expression statistics are used to rank genes by their propensity to be differentially expressed, which we then use to interpret the changes observed in a template experiment. This allows investigators to distinguish common DEGs from context specific ones in their results. We generate a z-score per gene to capture the relationship between a gene’s magnitude of change in the template experiment compared to the background distribution. In general, if a gene’s magnitude of change is larger than the mean change in the background distribution, then this gene is considered specific. However, the specificity threshold will depend on the experiment of interest and what additional contextual constraints being considered.

### Simulation-based approach identifies common DEGs that recapitulate curation-derived ones

Identifying common differential expression has been challenging because it requires extensive manual curation. We sought to compare the common DEGs identified by SOPHIE with those identified in a prior report. The prior study curated 2,456 human microarray datasets from the GPL570 (Affymetrix Human Genome U133 Plus 2.0 Array) platform to identify common DEGs.^6^ This study provided a list of genes ranked based on how frequently they were identified as differentially expressed across approximately 600 experiments, which we refer to as the Crow *et al.* results. We compared SOPHIE-predicted common DEGs using a VAE trained on the Crow *et al.* dataset with the results reported in Crow *et a*l. We calculated the percentile of genes by their median log_2_ fold change across the 25 simulated experiments. Comparing the gene percentiles from Crow *et al.* to our SOPHIE results revealed substantial concordance (Figure 1B; Spearman correlation coefficient at 0.591). There was also a significant (p-value<1e-16) over-representation of SOPHIE identified common DEGs within the common changes that Crow *et al.* identified. SOPHIE recapitulated the primary results of the curation-based approach for Crow *et al.* While Crow *et al.* relied on having a manually curated dataset, SOPHIE identified these genes in a more scalable and automated way, leveraging existing gene expression data to simulate a background set of experiments to use as a reference.

### SOPHIE finds common DEGs are consistent across contexts and platforms

We next examined whether or not common differentially expressed genes were consistent across training datasets and platforms. We applied SOPHIE using a different collection of microarray data that accompanied another prior report of commonly differentially expressed pathways.^7^ This second dataset we refer to as the Powers *et al.* results, which included 442 differential expression analyses (from 2,812 human microarray datasets) testing the response of small-molecule treatments in cancer cell lines. For this analysis, we selected an arbitrary template experiment (GSE11352 examined estradiol exposure in breast cancer cells^15^) to generate simulated experiments. We calculated differential expression statistics for each experiment and then calculated the percentile of genes by their median log_2_ fold change across the simulated experiments. We found concordance between SOPHIE-identified common DEGs using a VAE trained on Powers *et al*. and the results published in Crow *et al.* using Spearman correlation (Figure 2A). The concordance was particularly high for the genes in the highest and lowest percentiles, the most and least commonly differentially expressed genes respectively. Furthermore, there was a significant (p-value=1e-49) over-representation of SOPHIE identified common DEGs within the common changes that Crow *et al.* identified. While the two datasets used the same array platform to generate data, the datasets have different compositions – Crow *et al.* is a heterogenous mixture of different types of experiments while Power *et al.* is specifically cancer cell lines treated with small molecules. The consistency we observe in the common DEGs despite the differences in context demonstrates that many common DEGs are differentially expressed regardless of the context.

**Figure 2:**
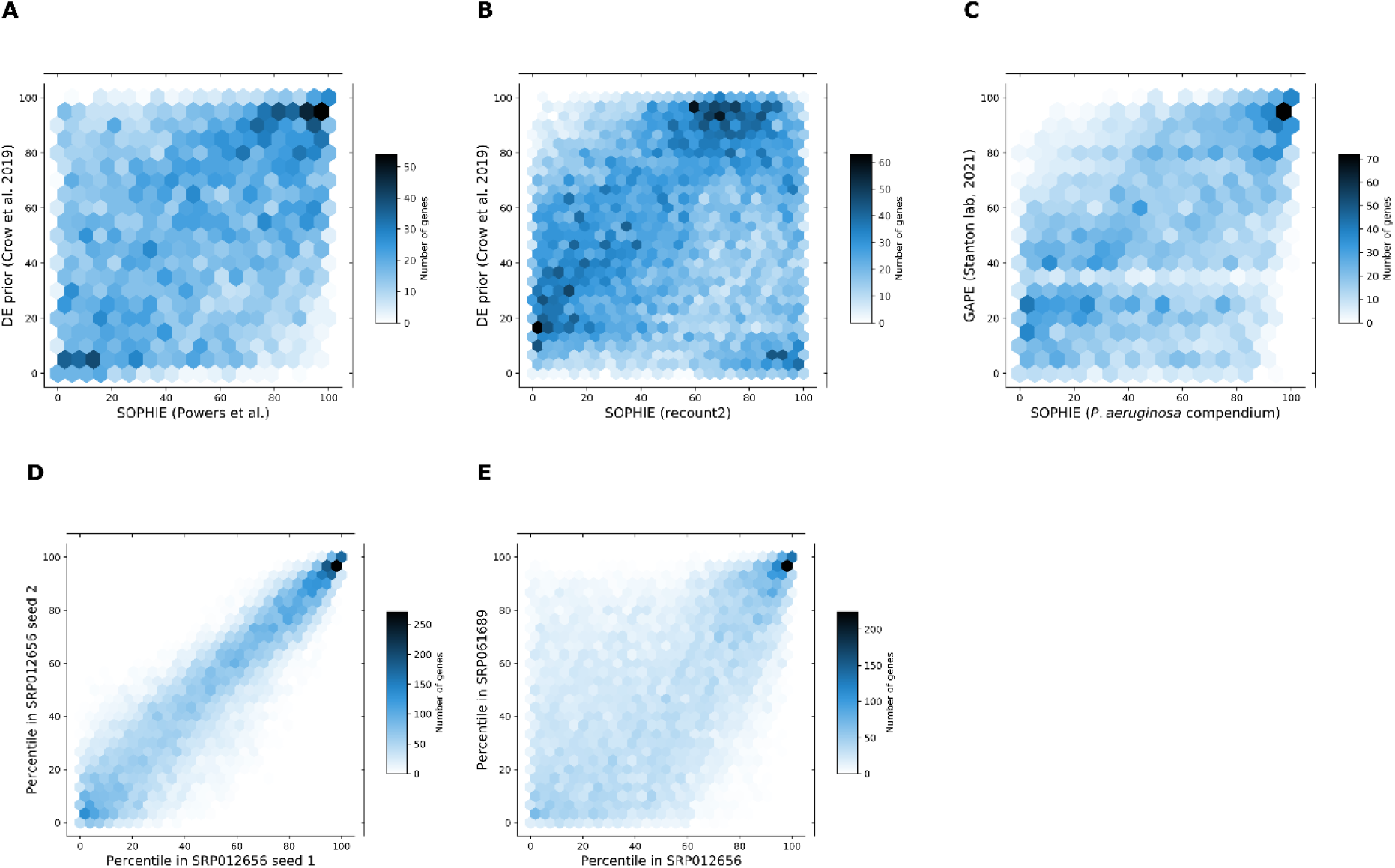
SOPHIE finds some common DEGs that are consistent across different platforms and contexts. A) Spearman correlation between gene percentiles using SOPHIE trained on Powers *et al.* (array) using GSE11352 as a template (x-axis) versus percentiles using manually curated experiments from Crow *et al.* (y-axis, same array platform but different context) with significant over-representation of SOPHIE common DEGs in Crow *et al.* common DEGs (p-value=1e-49). B) Spearman correlation between gene percentiles using SOPHIE trained on recount2 (RNA-seq) using SRP012656 as a template (x-axis) versus percentile using manually curated experiments from Crow *et al.* (y-axis, array) with significant over-representation of SOPHIE common DEGs in Crow *et al.* common DEGs (p-value=2e-15). SOPHIE can also easily extend to find common DEGs in different organisms. C) Spearman correlation between gene percentile using SOPHIE trained on the *P. aeruginosa* compendium (array) using E-GEOD-33245 as a template (x-axis) versus percentile using manually curated experiments from GAPE. (y-axis) with significant over-representation of SOPHIE common DEGs in GAPE common DEGs (p-value=1e-139). SOPHIE findings are robust. D) Spearman correlation (R^2^ = 0.907) between gene percentiles generated by SOPHIE using two runs of the same experiment (SRP012656) and E) Spearman correlation (R^2^=0.572) between gene percentiles generated by SOPHIE using two different template experiments (SRP012656 and SRP061689).

In general, transcriptome analysis approaches can be difficult to translate between different platforms (RNA-seq, microarray) and datasets. To demonstrate whether common DEGs were consistent across platforms, we applied SOPHIE using human RNA-seq data from recount2^12^. We selected an arbitrary template experiment from recount2 (SRP012656 examined non-small cell lung adenocarcinoma tumors^16^), simulated experiments and calculated differentially expressed genes using DESeq2. For this template experiment, primary non-small cell lung adenocarcinoma tumors were compared to adjacent normal tissues for 6 never-smoker Korean female patients. We again examined concordance compared to the common DEGs reported in Crow *et al* (Figure 2B). Despite the Crow *et al.* data being measured on microarrays while recount2 used an RNA-seq platform, we still found a significant (p-value= 2e-15) over-representation of SOPHIE-identified common DEGs shared with the Crow *et al.* analysis.

We also noticed a set of genes in the bottom right corner of Figure 2B with a high percentile score that were common DEGs in RNA-seq but not in Crow *et al.* We did not observe a corresponding set in the upper left corner, suggesting that RNA-seq captured the microarray-based common DEGs, but prior microarray-based reports lacked certain RNA-seq specific ones. This subset of genes was commonly differentially expressed in RNA-seq and not in array data, suggesting that platform differences underlie this effect. Some preliminary experiments showed that common DEGs identified specifically in the RNA-seq data tended to have a lower expression compared to those common DEGs identified using both the array and RNA-seq platform (Figure S1). The VAE, used by ponyo in the simulation step, appeared to artificially boost the expression of these RNA-seq-identified common DEGs so that they were found to be differentially expressed. Unlike the array data, the RNA-seq data has a larger variance and so the effects of the VAE are more pronounced, affecting genes in the outliers of the compendium distribution, which includes these RNA-seq identified commonly differentially expressed genes. In general, there was a consistent set of common DEGs found using two datasets that have similar contexts – they both contain a mixture of different types of experiments – but used different platforms. This consistency indicates that there are some common DEGs that are differentially expressed across different platforms.

Overall, using SOPHIE we found that there exists some common DEGs that are consistent across contexts and platforms – there is a set of frequently differentially expressed genes, regardless of context or platform (Figure S2).

### SOPHIE generalizes to other organisms

Finally, when we extended SOPHIE to a different organism, *P. aeruginosa*, we observed concordance (R^2^ = 0.449) between SOPHIE-generated percentiles compared to those generated using a manually curated dataset, GAPE (Figure 2C).^17^ GAPE contained a collection of 73 array experiments from the GPL84 platform. GAPE performed automatic group assignments of those experiments that were then manually verified by human curators. We then calculated the percentile for how frequently genes were differentially expressed across the 73 experiments. For this analysis, we selected the template experiment E-GEOD-33245, which examined different targets of the carbon catabolite control system^14^, to generate simulated experiments. We calculated differential expression statistics for each experiment and then calculated the percentile of genes by their median log_2_ fold change across the simulated experiments. We found a significant over-representation (p=1e-139) of SOPHIE identified common DEGs within the GAPE set of common DEGs. Again, without any curation, SOPHIE recapitulated the common findings reported in the GAPE dataset, which was generated using a manually curated approach. With our previous analysis using human data, the consistency found in these results demonstrate the generalizability of SOPHIE to other organisms like bacteria – with our SOPHIE approach we could easily switch out the human training dataset with a bacterial one.

### SOPHIE common findings are robust

Having shown that SOPHIE can recapitulate the commonly differentially expressed gene percentiles identified by two manually curated datasets (Crow *et al.* or GAPE) using a variety of input datasets, we next examined the robustness of these common patterns using a human compendium. We compared SOPHIE percentiles from different simulations using the same template experiment and found a very strong correlation (R^2^ = 0.907), especially for high and low percentile genes (Figure 2D). The genes in the middle percentiles are more sensitive to changes so the signal is less clear, but this is not unexpected with rank-based analysis in gene expression, where small changes near the middle of the distribution can produce large differences in rank. This noise is more pronounced when we compare the percentiles generated using two different template experiments (Figure 2E). Overall, we observe consistent common DEG percentiles across different template experiments (R^2^=0.572). SOPHIE common findings are robust to different runs and template experiments selected.

### Commonly differentially expressed pathways identified by SOPHIE recapitulate curation-derived ones

In addition to common DEGs, we also examined common differentially expressed pathways. While there is some variation between the ranking of common DEGs, grouping genes into pathways may find more robust common signals. For this analysis we used a set of common differentially expressed pathways reported by Powers *et al*. We calculated the percentile per pathway by how frequently enriched they were across the 442 experiments. Then, similar to the previous analyses, we applied SOPHIE, using the same Powers *et al.* data. We simulated 25 new experiments from the same template experiment used previously (GSE11352) and calculated differential expression statistics for each experiment. For this analysis, since we are focused on pathways, we then used GSEA^18^ to identify pathways enriched in differentially expressed genes. We compared the percentile of pathways determined using data simulated from SOPHIE with those we calculated based on the reported by Powers *et al.* and found strong concordance (R^2^= 0.65, Figure 3A). SOPHIE recapitulated the commonly enriched pathways reported in Powers et al, which used a manual curation approach.

**Figure 3:**
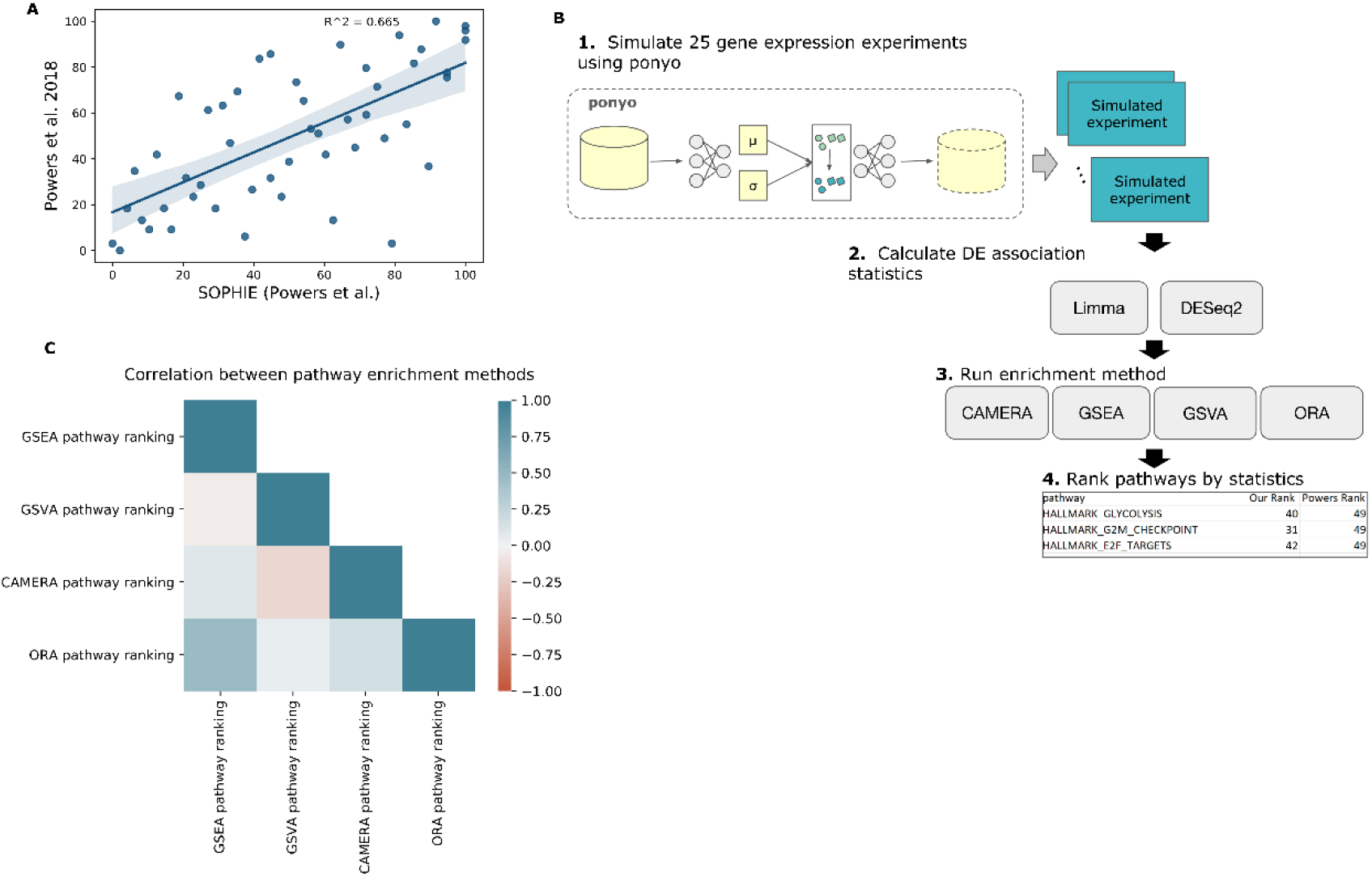
SOPHIE identifies the same commonly changed pathways previously found using manual curation. A) Correlation between pathway percentiles using our simulated method trained on Powers *et al.* compendium (x-axis) versus percentiles obtained from Powers *et al.* (y-axis). B) Workflow describing how the SOPHIE pipeline can be easily extended to plug in different enrichment methods. C) Correlation of pathway percentiles between different enrichment methods (GSEA, GSVA, CAMERA, ORA) using RNA-seq data.

SOPHIE can also be applied using other pathway analysis methods. We easily extended SOPHIE to use multiple different enrichment methods (Figure 3B) and examined the common findings. We selected 4 enrichment methods (GSEA, GSVA, CAMERA, ORA) from Geistlinger *et al.*^19^ We selected methods if 1) they could be applied to both RNA-seq and array data and 2) they covered a wide range of statistical performance measures including runtime, the number of gene sets found to be statistically significant and the type of method – self-contained versus competitive. Overall, the percentile of common pathways enriched varied between enrichment methods, likely due to the different assumptions and modeling procedures (Figure 3C, S3). Therefore, scientists will need to use a method-specific common correction approach. Similar to our analysis of common DEGs, compared to Powers *et al.*, SOPHIE can automatically identify commonly changed pathways. Additionally, SOPHIE can be easily customized to use different enrichment methods depending on the analysis.

### Common DEGs may correspond to hyperresponsive pathways

We next examined how the genes that are commonly differentially expressed are related to previously reported transcriptional patterns to gain insight into the role of these common DEGs. We identified common DEGs using recount2, which is a heterogeneous compendium of human gene expression data containing a range of different types of experiments and tissue types. The recount2 data was decomposed into latent variables (LV), representing gene expression modules, some of which were aligned with known curated pathways, in prior work.^20^ In these latent variables, genes had some weighted contribution, and we found that the median number of genes with non-zero weight was 2,824. We divided genes into a set of common DEGs, which were genes that were in the 60^th^ percentile and above in our recount2 analysis (Figure 2B), and all other genes. We found that the common DEGs had non-zero weight to roughly the same number of latent variables as non-common DEGs (other genes) (Figure 4A, p-value = 0.239 comparing the median between gene groups). However, common DEGs were found among the highest weights (the 98^th^ percentile and above for each latent variable) for fewer latent variables than other genes (Figure 4B, p-value=6e-119 comparing the median number of highly contributing genes between common DEGs with other genes). Taken together, these results suggest that common DEGs contribute to as many latent variables as other genes (i.e. have a non-zero weight), but common DEGs occur less frequently among the highest weight genes. Overall, the wide coverage across latent variables but lack of high weight contributions suggests that common DEGs across human experiments mainly contribute to a few pathways.

**Figure 4:**
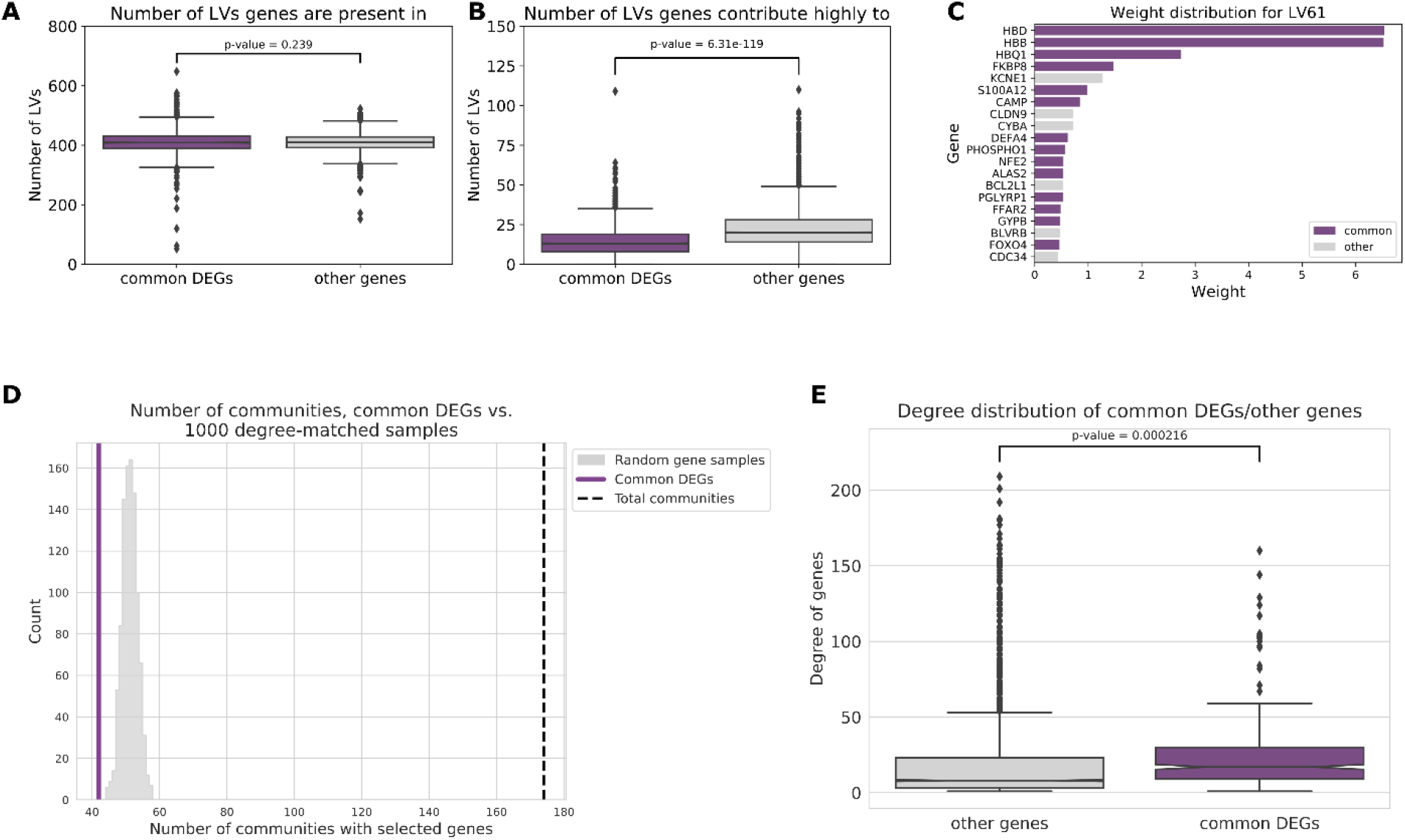
Common DEGs may contribute to a few hyperresponsive pathways. A) Number of human PLIER latent variables (LVs) common DEGs and other genes are present in (t-test p-value=0.239). B) Number of human PLIER latent variables common DEGs and other genes have a high weight score in (t-test p-value=6.31e-119). C) Distribution of top-weighted human genes in example LV61, which was found to contain a high proportion of high weight common DEGs. D) The number of communities with at least one commonly changed *P. aeruginosa* gene (purple) compared to the distribution of the number of communities with at least one non-commonly changed gene across 1000 samplings (grey) with the total number of communities marked by the black dashed line. E) Distribution of the degree of commonly changed *P. aeruginosa* genes (purple) compared to other genes (grey).

Given the small number of latent variables that common DEGs are high weight in, one possibility for why these genes were commonly changed might be related to membership in a few hyper-responsive pathways. Since these latent variables tend to be associated with particular biological processes, we tested if there were any latent variables, and thereby processes, that contained a large fraction of common DEGs. If there exist latent variables that were primarily composed of common DEGs, this might lend insight into the role of commonly differentially expressed genes. For this analysis, we ranked latent variables by the proportion of commonly shifted genes at the 98^th^ percentile and above. Overall, many of these latent variables were associated with immune responses, signaling, and metabolism. One example latent variable, that contained a high proportion of common DEGs compared to other genes (proportion of common DEGs > 0.5), was LV61 (Figure 4C, Supplementary Table S1). This latent variable included pathways related to immune response (Neutrophils), signaling (DMAP ERY2), and wound healing (megakaryocyte platelet production).

We performed a similar analysis to examine common patterns in *P. aeruginosa* data. Again, we leveraged an existing model. Tan *et al.* previously created a low dimensional representation of the *P. aeruginosa* compendium using a denoising autoencoder, called eADAGE, where some of the latent variables were found to be associated with KEGG pathways and other biological sources of variation.^21–23^ Using this existing eADAGE model, we created a gene-gene similarity network where the correlation within the eADAGE representation was used to connect genes. After performing a community detection analysis, we discovered that common DEGs, those genes with high concordance between SOPHIE and GAPE, tended to cluster in fewer communities compared to other genes (Figure 4D, Supplementary Table S2). Furthermore, common DEGs had a slightly higher median degree in the eADAGE similarity network compared to other genes (Figure 4E). These observations were consistent with an analysis that found a set of virulence-related transcriptional regulators that target multiple pathways.^24^ Together, these data suggest that, like the patterns we observed in the human dataset, there are relatively few communities that common DEGs changed genes contribute strongly to. These few communities containing common DEGs were highly connected to other communities, again suggesting that certain pathways may be particularly responsive to perturbations.

### SOPHIE-identified common DEGs involved in, but not specific to, the carbon catabolite repression system in P. aeruginosa

In general, differential expression analyses often aim to understand the genetic causes and downstream consequences of gene expression. However, using traditional p-values and log fold change criteria, such datasets often contain hundreds of genes, many of which are secondary to changes in the phenotype of interest. Using SOPHIE, we distinguish between common DEGs versus those that are specific to the context of the experiment. As a test case, we examined the common and specific genes generated using the template experiment E-GEOD-33245 which investigated the metabolic decision-making process known as carbon catabolite repression, that is important for *P. aeruginosa* pathogenicity^25^ (Figure 5A).

**Figure 5:**
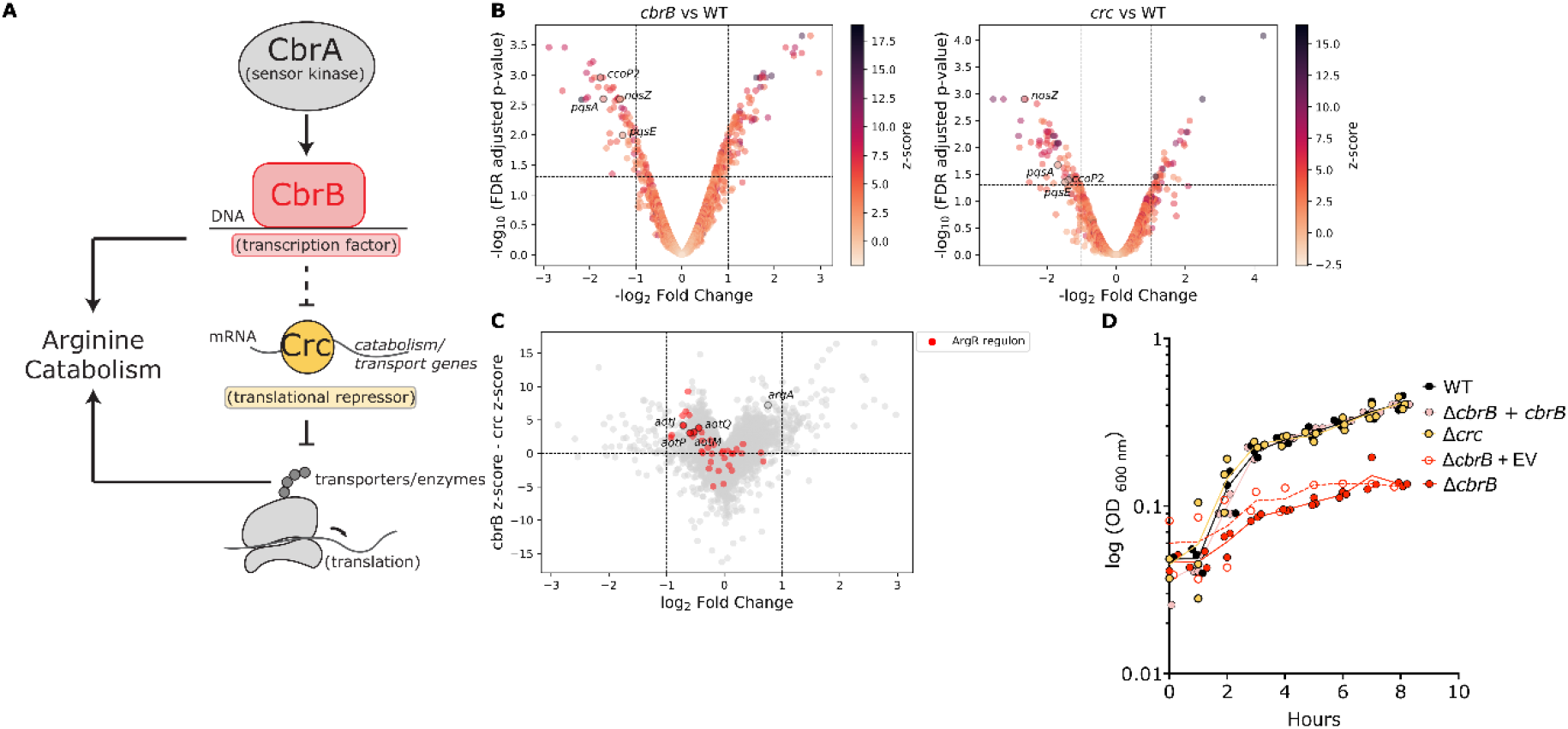
SOPHIE can identify genes with specific expression shifts in experiments. A) Model of CbrAB system. Volcano plot with log_2_ fold change versus adjusted p-values for B) WT vs *cbrB* mutant and WT vs *crc* mutant. The darker hue indicates a higher z-score and therefore higher specificity for the context being tested. C) Plot with log_2_ fold change in *cbrB* mutant context on the x-axis and difference in z-score in *cbrB* and *crc* mutant contexts on the y-axis. So changes that are specific to *cbrB* have positive y-values and changes specific to *crc* have negative y-values. D) Growth curves for *P. aeruginosa* in 10 mM arginine using WT (black), *cbrB* mutant (filled red), *cbrB* mutant with an empty expression vector (empty red), *cbrB* mutant with extrachromosomal complementation (pink), and *crc* mutant (yellow). Note: *cbrB* and *crc* were removed when plotting panels B and C.

To separate common and context specific DEGs, we used the z-score that compares the log_2_ fold change of a gene in a template experiment to the mean log_2_ fold change of that same gene across the background set of experiments. A low z-score indicated that there was no significant difference in how changed the gene was between the template versus background set and therefore these genes were predicted to be common DEGs.

Genes that had a low z-score, indicating a high likelihood of it being part of a common response, were differentially expressed in many experiments across the *P. aeruginosa* datasets: genes considered commonly differentially expressed by SOPHIE and GAPE accounted for a substantial fraction of differentially expressed genes in Δ*cbrB* and *Δcrc* comparisons respectively (Figure 5B). Both comparisons included the well-studied genes *pqsA, pqsE*, *nosZ*, and *ccoP2* as commonly differentially expressed. One differentially expressed gene in the Δ*crc* comparison with wildtype was *arcB,* an ornithine carbamoyltransferase involved in the arginine deiminase pathway that produces ornithine from arginine under low oxygen conditions. Based on SOPHIE analysis, this gene had a z-score of 1.09 to suggest it is a commonly differentially expressed gene. This assignment as a common DEG aligns well with the published GAPE analysis that found *arcB* to be differentially expressed in 40 out of the 73 annotated *P. aeruginosa* studies.

### SOPHIE identified arginine catabolism genes as specific to components in the carbon catabolite repression system

In addition to the identification of common DEGs, an orthogonal use of SOPHIE can be applied when analyzing experimental conditions that uncover unrecognized but specific genes of interest. In separating common and specific DEGs, SOPHIE can highlight those that show modest, but specific changes that would be missed by traditional DE analysis. This use is applicable to the carbon catabolite repression dataset (E-GEOD-33245) which included investigations into multiple genetic components of the same molecular pathway that collectively controls metabolic decision making. Ultimately, this pathway determines the order of metabolite consumption. This decision process depends on a complex molecular mechanism involving both transcriptional and translational regulation that results in both direct and indirect effects on the transcriptome respectively. A previous analysis by Sonnleitner *et al*.^14^ suggested that the production of catabolic enzymes and transporters is controlled by the translational co-repressor Crc (Figure 5A). In the presence of non-repressive carbon sources, the CbrA kinase promotes activity of the CbrB transcriptional regulator, which directly modulates levels of the small RNA *crcZ* among other transcripts. In turn, *crcZ* sequesters the Crc protein^26^ thereby enabling translation to occur.

We focused on the comparisons between WT and isogenic Δ*cbrB* and *Δcrc* mutants from E-GEOD-33245 and sought to identify transcriptional changes specific to one or the other regulator. In the absence of the transcription factor CbrB or the translational co-repressor Crc, 156 and 149 genes were differentially expressed (|log_2_FC| > 1, FDR-adj p-value < 0.05), respectively, relative to wild type. To select context-specific DEGs, we again used the z-score that compared the log_2_ fold change of a gene in a template experiment compared to the mean log_2_ fold change of that same gene across the background set of experiments, this time selecting for large z-scores. If a z-score was large, then the gene is more differentially expressed in the template experiment compared to the background set of experiments and therefore predicted to be specific to the template experiment. In our case, we selected genes that had a large z-score and that were specific in one condition versus the other, so our z-scores were not necessarily the largest overall. Depending on the use case, scientists will need to determine which z-scores are large enough given the contextual constraints to consider.

SOPHIE revealed genes involved in aerobic arginine metabolism (*argA*) and arginine transport (*aotJQMP)* changed by less than 2-fold in both samples. However, although CbrB and Crc are part of the same metabolic regulatory pathway, the specificity (high ranked z-score, Supplementary Table S3) was high in Δ*cbrB* but not Δ*crc*. Broadly, genes regulated by the arginine responsive regulator ArgR were more specific to deletion of *cbrB* than *crc* (Figure 5C, supplementary Table S3).^27^ We constructed *P. aeruginosa* strain PA14 mutants Δ*cbrB* and Δ*crc* and found that only Δ*cbrB* was defective for growth on arginine likely the result of defective transport or catabolism (Figure 5D). This result supports the model that arginine metabolism is specifically regulated by CbrB, consistent with published data by other studies^28, 29^, and highlights the utility of SOPHIE to drive the prioritization of genes for follow-up analysis of candidate differentially expressed genes. This method is particularly powerful for those genes that do not change very much but do so more than in the background simulated experiments (i.e. specific genes). It is appreciated that small expression changes can have biological significance, but we often choose not to pursue these genes because it is more difficult to study and follow low expression changes. However, SOPHIE provides strong confidence scores that highlight biologically important, but less studied genes for further analysis. By leveraging publicly available data, SOPHIE identified candidate specific genes. Independently, we experimentally validated that these genes played a specific role in the context of the template experiment. SOPHIE can therefore successfully predict biologically relevant gene targets that further our mechanistic understanding and drive future analyses.

## Discussion

We introduce an approach, SOPHIE, named after one of the main characters from Hayao Miyazaki’s animated film *Howl’s moving castle*. Sophie’s outward appearance as an old woman, despite being a young woman that has been cursed, demonstrates that initial observations can be misleading. This is the idea behind out approach, which allows users to identify specific gene expression signatures that can be masked by common background patterns.

SOPHIE automatically identified commonly differentially expressed genes and pathways using public gene expression compendia. SOPHIE returned consistent genes and pathways, by percentile, compared to previous results using both human^6, 7, 12^ and bacterial^22^ datasets. SOPHIE also found that many common DEGs were consistent across contexts and platforms. Furthermore, experimental validation confirmed a group of genes that SOPHIE predicted to show context-specific differential expression. In contrast to using a manually curated dataset, SOPHIE can be easily extended to generate a background distribution of experiments for any organism with public data available. These background experiments define a set of genes and pathways that are commonly changed across many different experimental conditions. These background sets of changes, provide context to individual experiments, highlighting specific gene expression changes and thus giving insight into mechanisms relevant to specific contexts including disease conditions.

Compared to prior work using manually curated datasets, which required laborious manual grouping^6, 7, 17^, SOPHIE demonstrates consistent results but using an automated process. In short, SOPHIE identifies the same common patterns but in a fast and scalable way. However, there was a subset of genes that were specifically differentially expressed using SOPHIE but not found using the manually curated background. In one case, SOPHIE is using RNA-seq while the manually curated data is based on hybridization technology (microarray). Some initial experiments showed that this inconsistency is likely due to platform differences and how the VAE handled these two different data types. Overall, SOPHIE results are consistent with previous findings regardless of platform, but we also identified differences that might indicate there exists a hierarchy of common changes depending on the platform.

Building on the discovery of these common signals, we also examined the potential role of these commonly differentially expressed genes. These common DEGs appear to contribute to a small number of hyperresponsive pathways (Figure 4). This supports the observation that genes found to be differentially expressed across different contexts may not be informative about the experimental manipulation of interest. Therefore, considering specificity can be complementary to using log fold change activity to study biological processes.

SOPHIE is a general approach relying on generative neural networks. Depending on the data type, there likely exists some optimal neural network architecture that preserves the underlying structure in the data. In our case, we examine SOPHIE with a VAE. VAEs can inappropriately reduce the variance in the data due to the normality assumption^30^, potentially affecting the number of DEGs. However, while this limitation is known, Lee et al.^8^ demonstrated that VAEs can still produce realistic experiments in this context. Based on this limitation, we used SOPHIE with percentile ranks, aligning with prior work from Crow et al.^6^, instead of raw values to identify common DEGs. While using a VAE was successful at allowing us to identify common DEGs in our SOPHIE framework, other generative neural networks may be superior and future work is needed to optimize and assess different types of generative neural networks to determine what model is most appropriate for a given dataset, data type, or measurement platform.

One limitation is that our template experiments are comprised of two conditions, but there are many different types of experiments (e.g. time course). To determine if common DEGs vary based on experiment design, we would need to curate more experiments testing different experimental designs and determine how to group samples to perform a differential expression analysis or develop a new metric to define how many genes change. Another limitation to our study is that ponyo uses a random linear shift to simulate experiments. While this linear shift is using a location drawn from the known distribution of gene expression data, this shift currently doesn’t allow us to vary or shift along certain axes, such as tissue type or drug. If ponyo could be extended to simulate background experiments along a specific axis, like tissue type or drug. To ask if there are different sets of common DEGs that come up as we vary along specific axes, we would need to have a deeper understanding of the structure of the latent space and what is being captured. These questions can help lead to an improved understanding of common signals and the type of correction that might be needed.

SOPHIE is a powerful approach that can be used to drive how we study mechanisms underlying different cellular states and diseases. With SOPHIE, we can identify common DEGs that might be useful for diagnostic^31^ and detection^32^ purposes. We can also identify specific signals that point to possible treatment options^33^. In general, studies trying to uncover these genetic mechanisms tend to focus on prominent biological signals – those genes that are strongly differentially expressed. However, with SOPHIE we can start to glean information about those genes that are subtle but specifically relevant to the biology in question. Overall, SOPHIE is a practice that can complement existing traditional analyses to separate specific versus common differentially expressed genes and pathways. These context-specific genes and pathways include both subtle changes that are largely unexplored and prominent changes that might point to areas of treatment and biomarker development. In general, SOPHIE can easily be applied across a range of different datasets to help drive discovery and further understanding of mechanisms.

The best way to use SOPHIE in practice will depend on the scientific question and the ease with which leads can be validated. The software associated with this paper is available on github (https://github.com/greenelab/generic-expression-patterns) and users can modify the notebooks for their own analysis following the instructions in the README file.

## Methods

### Gene expression datasets

We used four complementary gene expression compendia in this work. Three were sets of assays of human samples, two via microarray and the other via RNA-seq profiling. The fourth was a collection from the microbe *Pseudomonas aeruginosa*.

The first human compendium that we used contains gene expression data from Crow *et al.*^6^ We downloaded the dataset from Gemma on (March 20, 2021). Gemma contains public gene expression data primarily from GEO. Samples were selected using the GEO accession number provided in the supplementary material (“Exterinal.ID” column in Dataset S1). These samples were measured on the GPL570 (Affymetrix Human Genome U133 Plus 2.0 Array) platform, testing at least one condition and reporting at least one differentially expressed gene. Samples were processed using the *rma* library to convert probe intensity values from the *.cel* files to log_2_ base gene expression measurements, and these gene expression values were then log_10_ transformed to account for the large spread of the data and then normalized to 0-1 range per gene. We also had to remove a subset of genes and samples that contained NaNs, where the data was not available. This resulted in an expression matrix that contains 7,130 genes and 32,082 samples.

The first human compendium that we used contains gene expression data from Powers *et al.*^7^ We downloaded the dataset from synapse on (October 7, 2020). This dataset contains samples from the GEO measured on Affymetrix Human Genome U133 Plus 2.0 Array. Samples were selected based on the following criteria: having at least 2 replicates per condition and containing a vehicle control. The dataset included 442 experiments testing the response of small-molecule treatments in cancer cell lines. Samples were processed using the *rma* library to convert probe intensity values from the *.cel* files to log_2_ base gene expression measurements, and these gene expression values were then normalized to 0-1 range per gene. This resulted in an expression matrix that contains 6,763 genes and 2,410 samples.

The second human compendium that we used includes human RNA-seq data from recount2.^12^ We downloaded all SRA data in recount2 as RangedSummarizedExperiment (RSE) objects for each project id using the recount library in Bioconductor (version 1.12.0). Raw reads were mapped to genes using Rail-RNA^34^, which includes exon-exon splice junctions. Each RSE contained counts summarized at the gene level using the Gencode v25 (GRCh38.p7, CHR) annotation provided by Gencode.^35^ These RSE objects include coverage counts as opposed to read counts, so we applied the scale_counts function to scale by sample coverage (average number of reads mapped per nucleotide). The compendium contained 49,651 samples with measurements for 58,129 genes. Our goal was to compare percentiles with ones provided by Crow *et al.*^1^, which required us to map the ensembl gene ids in recount2 to HGNC symbols. We used the intersection of genes between the recount2 and Crow *et al.* sets. This resulted in a gene expression matrix of 49,651 samples and 17,755 genes. We then normalized gene expression values to a 0-1 range per gene. This recount2 compendium contained a heterogeneous set of gene expression experiments – 31 tissue types (i.e. blood, lung), 57 cell types (i.e. stem, HeLa), multiple experimental designs (i.e. case-control, time-series).

The last compendium contained *P. aeruginosa* gene expression data that was collected and processed as described in Lee *et al.*^8^ The dataset was originally downloaded from the ADAGE^22^ GitHub repository (https://github.com/greenelab/adage/tree/master/Data_collection_processing). Raw microarray data (measured on the release of the GeneChip *P. aeruginosa* genome array and the time of data freeze in 2014) were downloaded as .cel files. Then *rma* was used to convert probe intensity values from the *.cel* files to log_2_ base gene expression measurements. These gene expression values were then normalized to 0-1 range per gene. The resulting matrix contained 989 samples and 5,549 genes that represent a wide range of gene expression patterns including characterization of clinical isolates from cystic fibrosis infections, differences between mutant versus WT, response to antibiotic treatment, microbial interactions, and the adaptation from water to GI tract infection.

#### SOPHIE: Specific cOntext Pattern Highlighting In ExpressionSimulate gene expression experiments using ponyo

Our simulation applied the experiment-level simulation approach from Lee *et al.*^8^ The configuration of the VAE we used was the same as in this previous publication – 2,500 features in the hidden layer and 30 latent space features. Each layer used a rectified linear unit (ReLU) activation function to combine weights from the previous layer. We performed a 75:25 split of the data for training and validation. The hyperparameters were manually adjusted based on a visual inspection of the validation loss outputs. Our optimal hyperparameter settings were: learning rate of 0.001, a batch size of 10, warmups set to 0.01. We trained 3 VAE models using Crow *et al.* (10 epochs), recount2 (40 epochs), Powers *et al.* (40 epochs), and the *P. aeruginosa* (100 epochs) compendia.

We selected a template experiment from our compendium (SRP012656 from recount2, GSE10281 from Crow *et al.*, GSE11352 from Powers *et al.*, and E-GEOD-33245 from *P. aeruginosa*). For the most part, the selected template experiments are assumed to come from a similar distribution as our background compendium. We simulated a new experiment by linearly shifting the selected template experiment to a new location in the latent space. This new location was randomly sampled from the distribution of the low dimensional representation of the trained gene expression compendium. The vector that connects the template experiment and the new location was added to the template experiment to create a new simulated experiment. This process was repeated 25 times to create 25 simulated experiments based on the single template experiment. In general, we found that downstream statistical results were robust to different numbers of simulated experiments so we used 25 experiments to compromise on the runtime of the downstream analyses (Figure S4).

#### Differential expression analysis

For the recount2 compendium we used the DESeq module in the DESeq2 library^36^ to calculate differential expression values for each gene comparing the two different conditions in the selected template experiment (SRP012656). The template experiment contained primary non-small cell lung adenocarcinoma tumors and adjacent normal tissues of 6 never-smoker Korean female patients. The differential expression analysis compared tumor vs normal. Following a similar procedure for the array-based datasets (the Crow *et al.* compendium, the Powers *et al.* compendium and *P. aeruginosa* compendium) we used the eBayes module in the limma library^37^ to calculate differential gene expression values for each gene. The output statistics include log_2_ fold change between the two conditions tested and p-values adjusted by Benjamini-Hochberg’s method to control for false discovery rate (FDR). The template experiment we used for the Crow *et al.* compendium is GSE10281, which examined the expression profiles of breast cancer cells treated with Letrozole. The template experiment we used for the Powers *et al.* compendium is GSE11352, which examined the transcriptional response of MCF7 breast cancer cells to estradiol treatment. So the differential expression analysis compared samples untreated versus treated. The template experiment we used to the *P. aeruginosa* compendium is E-GEOD-33245, contained multiple comparisons examining the CbrAB system. The two we focused on for our analysis compared WT vs *cbrB* and *crc* mutants in LB media.

For the *P. aeruginosa* experiment, differentially expressed genes were those with FDR adjusted cutoff (using Benjamini-Hochberg correction) < 0.05 and log_2_ absolute value fold-change >1, which are thresholds frequently used in practice.

#### Calculate specificity of each gene (z-score)

Using the association statistics from the differential expression analysis, we calculated a score to indicate if a gene was specifically differentially expressed in the template experiment. We calculated a z-score for each gene using the following formula:

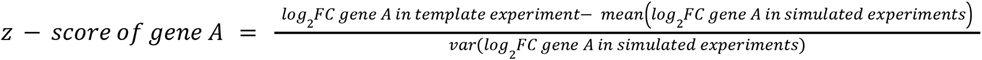

Higher z-scores indicate a gene is specifically differentially expressed in the template experiment in reference to the null set of experiments (i.e. 25 simulated experiments). This z-score is meant to guide scientists to select genes of interest. These genes could be the most specific gene (i.e. the genes with the highest z-scores) or it may be specific genes but those that follow other additional constraints and so the z-scores aren’t necessarily the highest.

### Enrichment analysis (EA)

The goal of EA is to detect coordinated changes in prespecified sets of related genes (i.e. those genes in the same pathway or share the same GO term).

Our primary method was GSEA, for which we used the fgsea module from the fgsea library.^18, 38^ The method first ranks all genes based on the DE association statistics. In this case, we used the log_2_ fold change. An enrichment score (ES) is defined as the maximum distance from the middle of the ranked list. Thus, the enrichment score indicates whether the genes contained in a gene set are clustered towards the beginning or the end of the ranked list (indicating a correlation with the change in expression). The statistical significance of the ES is estimated by a phenotypic-based permutation test to produce a null distribution for the ES (i.e. scores based on permuted phenotype). Each pathway was output with statistics including a Benjamini-Hochberg adjusted p-value. The pathways used in this analysis were the Hallmark pathways for the Powers *et al.* compendium

Other methods we used included: Gene Set Variation Analysis (GSVA)^39^, Correlation Adjusted Mean Rank gene set test (CAMERA)^40^, and Over-Representation Analysis (ORA). GSVA is a self-contained gene set test that estimates the variation of gene set enrichment over the samples independent of any class label. We used the gsva function from the gsva library. CAMERA is a competitive gene set test that performs the same rank-based test procedure as GSEA but also estimates the correlation between genes instead of treating genes independently. For CAMERA, we used the camera function that is part of the limma library.^41^ Last, ORA is a method that uses the hypergeometric test to determine if there a significant over-representation of a pathway in the selected set of DEGs. Here we used the clusterProfiler^42^ library but there are multiple options for this analysis.

### Comparison of gene percentiles

We wanted to compare the percentile of human genes identified using SOPHIE (trained on Crow *et al.*, Powers *et al.* and recount2 datasets) with the percentile found from *Crow et al.*, which identified a set of genes as common DEGs based on how frequently they were found to be DE across 635 manually curated experiments. In their paper, they ranked genes as 0 if they were not commonly DE and 1 if there were commonly DE. Our genes were ranked from 1 to 17,754 based on their median absolute log_2_ fold change value across the 25 simulated experiments. We linearly scaled the gene ranks to be a percentile from 0 to 100. Finally, we applied Spearman correlation to compare the percentile for each gene (Figure 1B, 2A, 2B).

We performed this same correlation analysis comparing SOPHIE trained on the *P. aeruginosa* compendium with percentiles generated from the GAPE project from the Stanton lab (https://github.com/DartmouthStantonLab/GAPE).^17^ The GAPE dataset contained ANOVA statistics generated for 73 *P. aeruginosa* microarray experiments using the Affymetrix platform GPL84. We downloaded the differential expression statistics for 73 array experiments from the associated repository (https://github.com/DartmouthStantonLab/GAPE/blob/main/Pa_GPL84_refine_ANOVA_List_unzip.rds). For each experiment, we identified differentially expressed genes using log_2_ fold change > 1 and FDR < 0.05. We then calculated the percentile per gene based on the proportion that they were found to be differentially expressed. We compared these GAPE percentiles against those found by SOPHIE (Figure 2C).

We also compared percentiles of genes amongst two SOPHIE-generated results. This included comparing percentiles generated from two SOPHIE runs using the same template experiment (Figure 2D) and SOPHIE generated for two different template experiments (Figure 2E).

### Comparison of pathway percentiles

We wanted to compare the percentile of pathways identified using SOPHIE (trained on Powers *et al.,* Crow *et al.*, and recount2 datasets) with the percentile based on the Powers *et al.* data. There was no pathway ranking provided in the publication, so we defined a reference ranking by calculating the fraction of the 442 experiments that a given pathway was found to be significant (FDR corrected p-value using Benjamini-Hochberg method <0.05) and used these rank pathways and then converted the ranking to a percentile as described above. We used the Hallmarks_qvalues_GSEAPreranked.csv file from https://www.synapse.org/#!Synapse:syn11806255. The file contains the q-values for the test: given the enrichment score (ES) of the experiment is significant compared to the null distribution of enrichment scores, where the null set is generated from permuted gene sets. Our percentile is based on the median Benjamini-Hochberg adjusted p-value across the simulated experiments. We compared our percentile versus the reference percentile using the Spearman correlation. We only show the comparison of SOPHIE trained on Powers *et al.,* but not Crow *et al.*, or recount2.

### Latent variable analysis

The goal of this analysis was to examine why genes were found to be commonly differentially expressed – we sought to answer the question: are common DEGs found in more Pathway-Level Information ExtractoR (PLIER) latent variables (LV)^20^ compared to specific genes? The PLIER model performed a matrix factorization of the same recount2 gene expression data to get two matrices: loadings (Z) and latent matrix (B). The loadings (Z) were constrained to aligned with curated pathways and gene sets specified by prior knowledge to ensure that some but not all latent variables capture known biology. For this analysis, we focused on the Z matrix, which is a weight matrix that has dimensions 6,750 genes by 987 LV. For this analysis, common DEGs were above the 60^th^ percentile (approximately the top 40% of genes were selected based on the distribution seen in Figure 4B) using the SOPHIE trained on recount2. We calculated the coverage of common DEGs versus other genes across these PLIER latent variables. For each gene we calculated two values: 1) how many LVs the gene was present in (i.e. has a nonzero weight value according to the Z matrix), 2) how many LVs the gene was high weight in, using the 98^th^ quantile for the LV distribution as the threshold.

### Network analysis

In order to examine associations between common differentially expressed genes and pathways or functional modules in *P. aeruginosa*, we constructed a network of gene-gene interactions. Nodes in this network represent *P. aeruginosa* genes, and edges represent correlations between the eADAGE weight vectors of the two genes they connect. We constructed the network using the ADAGEpath R package, described in more detail in the associated manuscript.^22^ To form the final network, we removed all edges (correlations) with a value between -0.5 and 0.5, and took the absolute value of the remaining edges (so negative edge weights became positive).

There are many existing methods to partition a network into well-connected, non-overlapping subnetworks, often referred to as communities. Using our gene similarity network, we sought to answer the question: Do common DEGs tend to occupy fewer network communities than a similar set of random genes, or do they tend to spread out across comparatively many communities? We chose two representative methods to divide the network into communities: (1) the Louvain method^43^, as implemented in the *python-igraph* package^44^, and (2) the “planted partition” model^45^ (data not shown), as implemented in the *graph-tool* Python package^45^. In order to make a meaningful comparison between common and non-common DEGs, we sampled an equal number of both gene categories. This meant that the non-common DEGs were approximately degree-matched with the common DEGs (i.e., for each commonly changed gene we sampled a specific differentially expressed gene with approximately the same network degree). We performed this sampling procedure 1000 times. We then counted the number of communities containing at least one commonly changed gene and compared this count to the distribution across the 1000 samples of the number of communities containing at least one sampled non-commonly changed gene.

In addition, we used the same eADAGE gene similarity network to compute several metrics describing individual network nodes, which we then compared between common and non-common DEGs. For both sets of genes, we calculated: (1) node degree, (2) edge weight, (3) betweenness centrality^46^ (4) PageRank centrality^47^. For each of these metrics, we used the implementations in the *graph-tool* Python package. In contrast to the other metrics, betweenness centrality treats edge weights as “costs” (lower = better, as opposed to correlation or similarity measures where higher = better), so for the betweenness centrality calculation we transformed all edge weights by setting edge cost = 1 - correlation.

### Strain Construction

Plasmids for making in-frame deletions of *cbrB* and *crc* were made using a *Saccharomyces cerevisiae* recombination technique previously described.^48^ The arabinose-inducible *cbrB* expression vector was made using Gibson cloning.^49^ All plasmids were sequenced at the Molecular Biology Core at the Geisel School of Medicine at Dartmouth and maintained in *E. coli*. In frame-deletions constructs were introduced into *P. aeruginosa* by conjugation via S17/lambda pir *E. coli.* Merodiploids were selected by drug resistance and double recombinants were obtained using sucrose counter-selection and genotype screening by PCR. The *cbrB* and empty expression vectors were introduced into *P. aeruginosa* by electroporation and selected by drug resistance.

### P. aeruginosa experiment

Bacteria were maintained on LB (lysogeny broth) with 1.5% agar. For strains harboring expression plasmids, 300 ug/mL Carbenicillin or 60 ug/mL Gentamycin was added. Yeast strains for cloning were maintained on YPD (yeast peptone dextrose) with 2% agar. Planktonic cultures (5 mL) were grown on roller drums at 37° from single colonies for 16 h in LB (under antibiotic selection for the appropriate strains). The 16 h LB cultures were normalized to OD_600 nm_ = 1 in 2 mL, and a 250 µL aliquot of the normalized culture was used to inoculate three 5 mL cultures of M63 medium containing 10 mM arginine as a sole carbon source under inducing conditions (0.2% arabinose) for a starting OD_600 nm_ = 0.05. Inoculated cultures were grown at 37° C on the roller drum and cellular density (OD_600 nm_) was monitored using a Spec20 every hour for 8 hours. Each data point is representative of the average of the 3 replicates per day for 3 independent days.

### Software

All scripts used in these analyses are available in the GitHub repository (https://github.com/greenelab/generic-expression-patterns) under an open-source license to facilitate reproducibility of these findings (BSD 3-Clause). We will archive this repository upon manuscript acceptance to Zenodo or a similar repository, and add the citation and persistent identifier here. The repository’s structure is described in the Readme file. The notebooks that perform the validation experiment for common DEGs and pathways can be found in “human_general_array_analysis” (SOPHIE trained on Crow *et al.*), “human_general_analysis” (SOPHIE trained on recount2), “human_cancer_analysis” (SOPHIE trained on Powers *et al.*), and “pseudomonas_analysis” (SOPHIE trained on the *P. aeruginosa* compendium) directories. The notebooks that explore why genes are commonly differentially expressed can be found in “LV_analysis” directory. The notebooks for the network analysis can be found in the “network_analysis” directory. All supporting functions to run these notebooks can be found in “generic_expression_patterns_modules” directory. The virtual environment was managed using conda (version 4.6.12), and the required libraries and packages are defined in the environment.yml file. Additionally, scripts to simulate gene expression experiments using the latent space shifting approach are available as a separate module, called *ponyo*, and can be installed from PyPi (https://github.com/greenelab/ponyo). The Readme file describes how users can re-run the analyses associated with this manuscript or analyze their own data using this method. An example of how to apply SOPHIE to a new dataset can be found in “new_experiment” directory. All simulations were run on a CPU.

## Acknowledgements

The authors would like to thank David Nicholson, Ben Heil, and Milton Pividori for reviewing the software associated with this work and providing valuable feedback. We would like to thank Ben Heil for coming up with the name of this method. This work was supported by grants from the Gordon and Betty Moore Foundation (GBMF4552 to CSG), the National Institutes of Health (R01 HG010067 to CSG, R01 CA237170 to CSG, U01 CA231978 to JCC), and the Cystic Fibrosis Foundation (HOGAN19G0 to DAH; GREENE21G0 to DAH and CSG). Support for the project was also provided by DartCF at the Geisel School of Medicine at Dartmouth to DAH, which is supported by NIH NIDDK grant P30 DK117469; the Cystic Fibrosis Foundation’s Research Development Program (CF RDP) under award STANTO19R0 and bioMT through NIH NIGMS grant P20 GM113132. The Gordon and Betty Moore Foundation (https://www.moore.org/), National Institutes of Health (https://www.nih.gov/) and Cystic Fibrosis Foundation (https://cff.org/), which supported this work, are available online at the linked locations. Finally we would like to thank Sabbi Lall from Live Sciences Editors for providing editing advice.

## Author Contributions

AJL: Formal analysis, investigation, methodology, project administration, software, validation, visualization, writing original, writing review

DLM: Formal analysis, investigation, visualization, writing original, writing review

JC: Formal analysis, investigation, software, writing review

DH: Software, writing review

RKP: Resources, writing review

GD: Writing review

JCC: Conceptualization, funding acquisition, supervision, writing review

DAH: Conceptualization, funding acquisition, supervision, writing review

CSG: Conceptualization, funding acquisition, supervision, writing original, writing review

---

**Figure S1:**
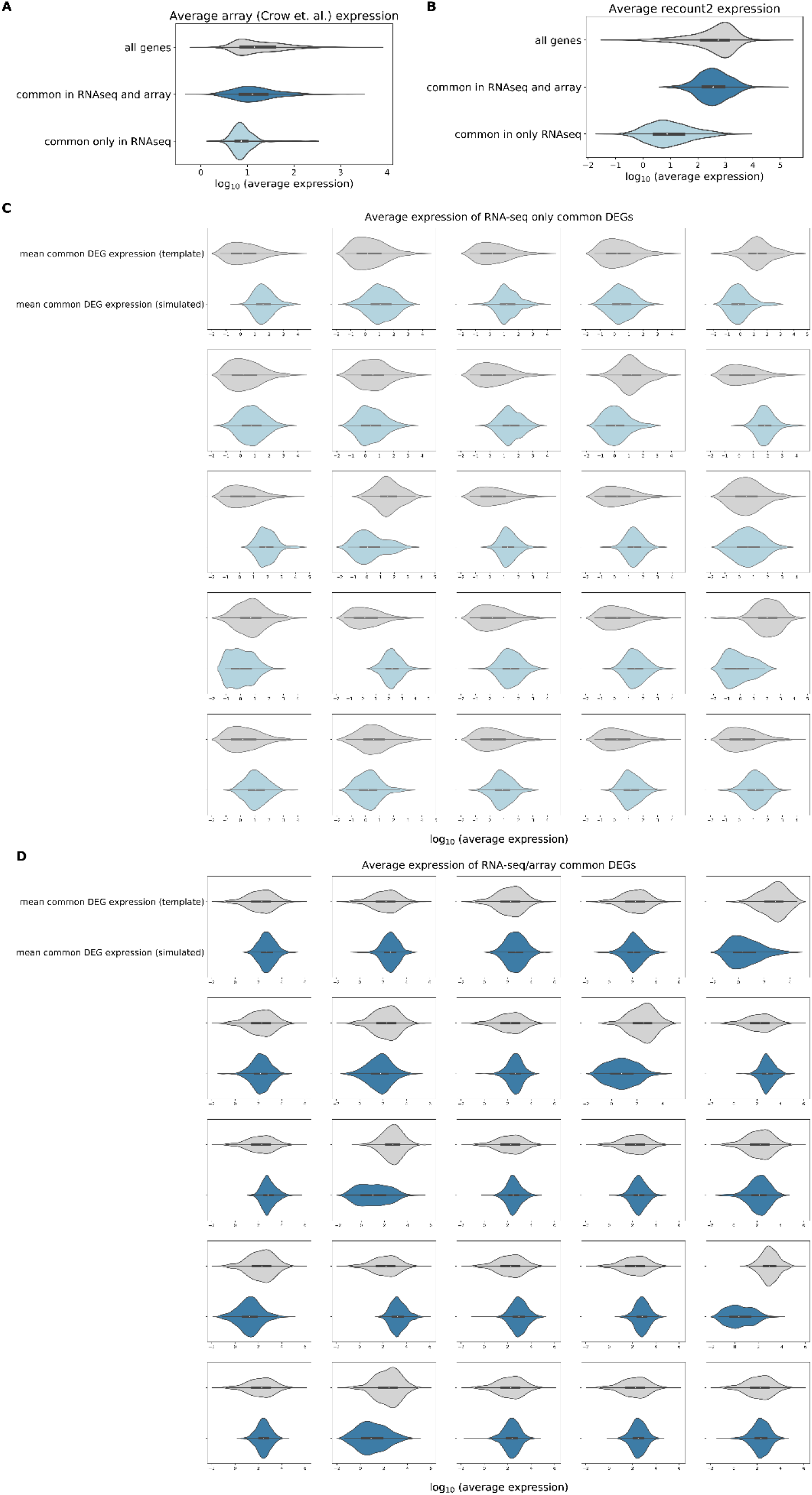
Common DEGs found in RNA-seq but not array data indicate platform-specific shifts. A) Average gene expression for all genes in Crow *et al.* array dataset (grey), genes commonly found to be changed in both RNA-seq using SOPHIE and array dataset using Crow *et al.* (dark blue), genes commonly found to be differentially expressed only in RNA-seq dataset (light blue). B) Average gene expression for all genes in recount2 RNA-seq dataset (grey), genes commonly found to be differentially expressed in both RNA-seq using SOPHIE and array dataset using Crow *et al.* (dark blue), genes commonly found to be differentially expressed only in RNA-seq dataset (light blue). C) Average gene expression of genes commonly found to be differentially expressed only in RNA-seq dataset in template experiment (grey) compared to simulated experiment (light blue). D) Average gene expression of genes commonly found to be shifted in both RNA-seq and array datasets in template experiment (grey) compared to simulated experiment (dark blue).

**Figure S2:**
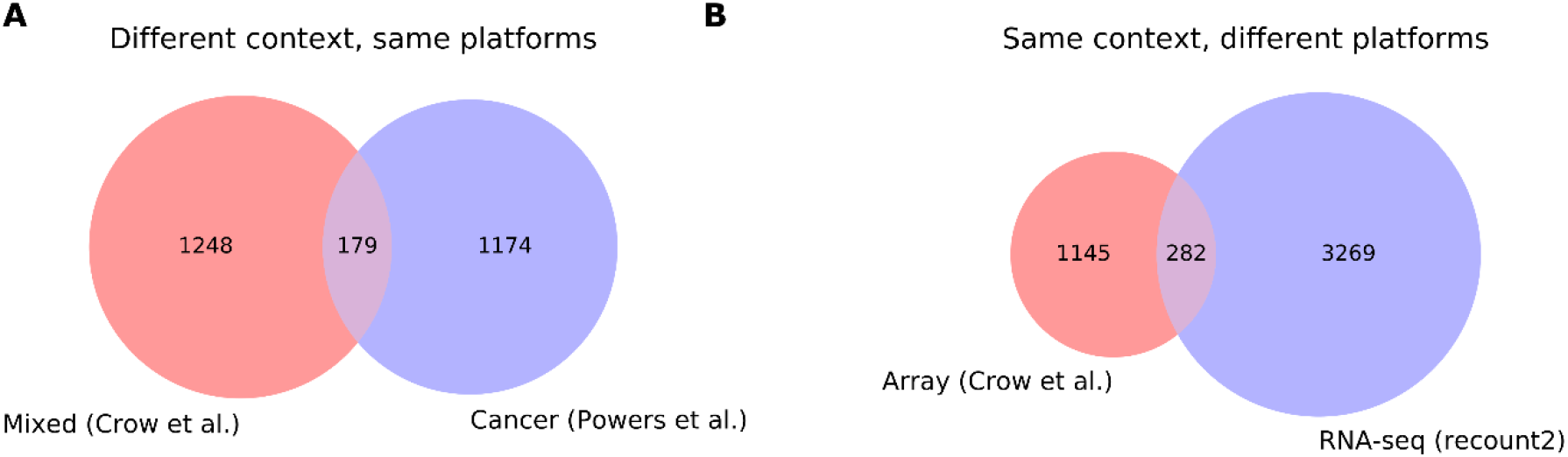
Consistency of common DEGs across context and compendia. A) Overlap of the top 20% most commonly changed genes identified using a heterogeneous compendium (Crow et al.) compared to using a cancer-specific compendium (Powers et al.). B) Overlap of the top 20% most commonly changed genes identified using a compendium composed of experiments measured on array (Crow et al.) compared to one measured on RNA-seq (recount2).

**Figure S3:**
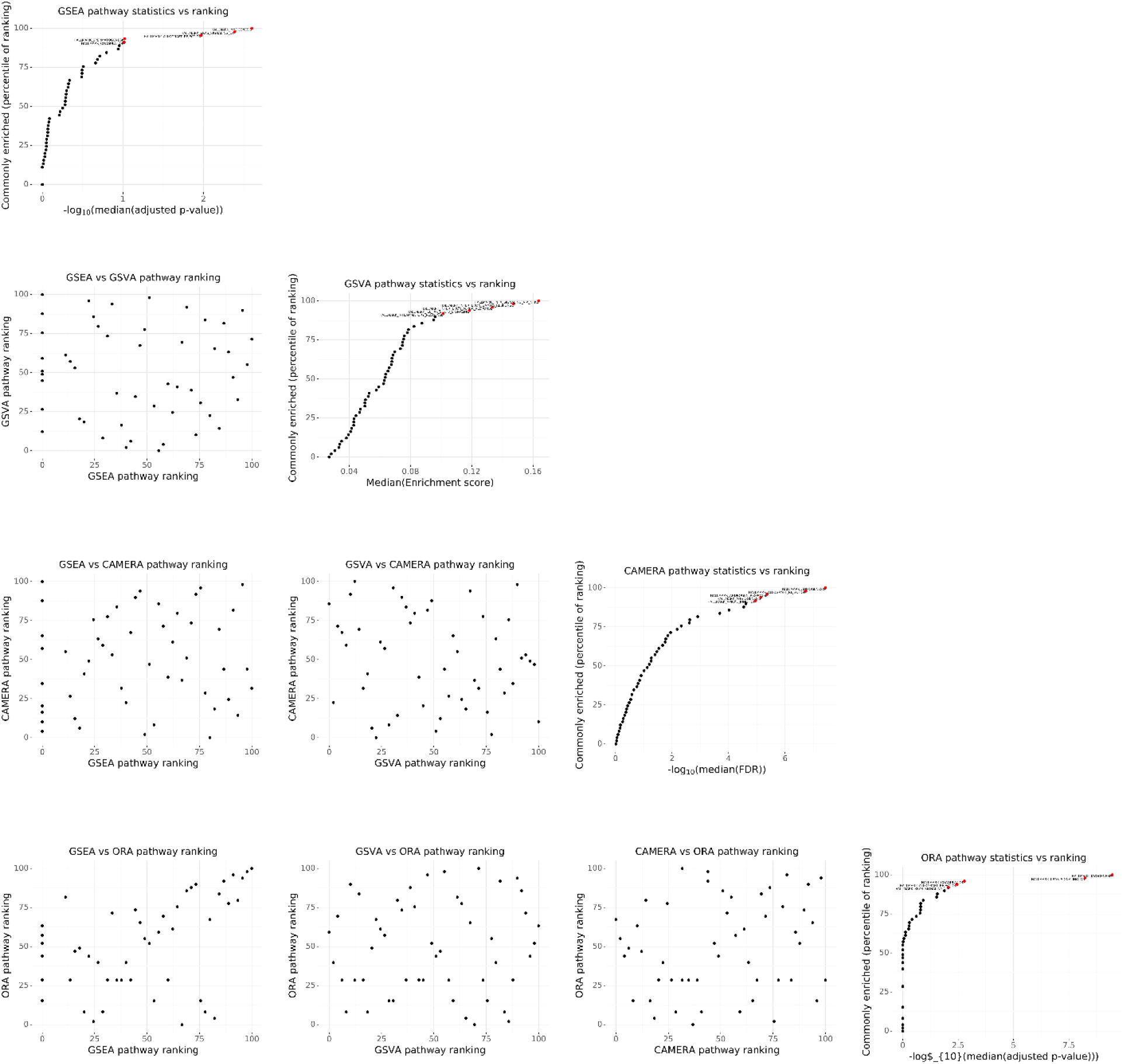
Different pathway enrichment methods will find different commonly enriched pathways. Scatterplot showing the correlation of pathway percentiles between different enrichment methods (GSEA, GSVA, CAMERA, ORA) using RNA-seq data.

**Figure S4:**
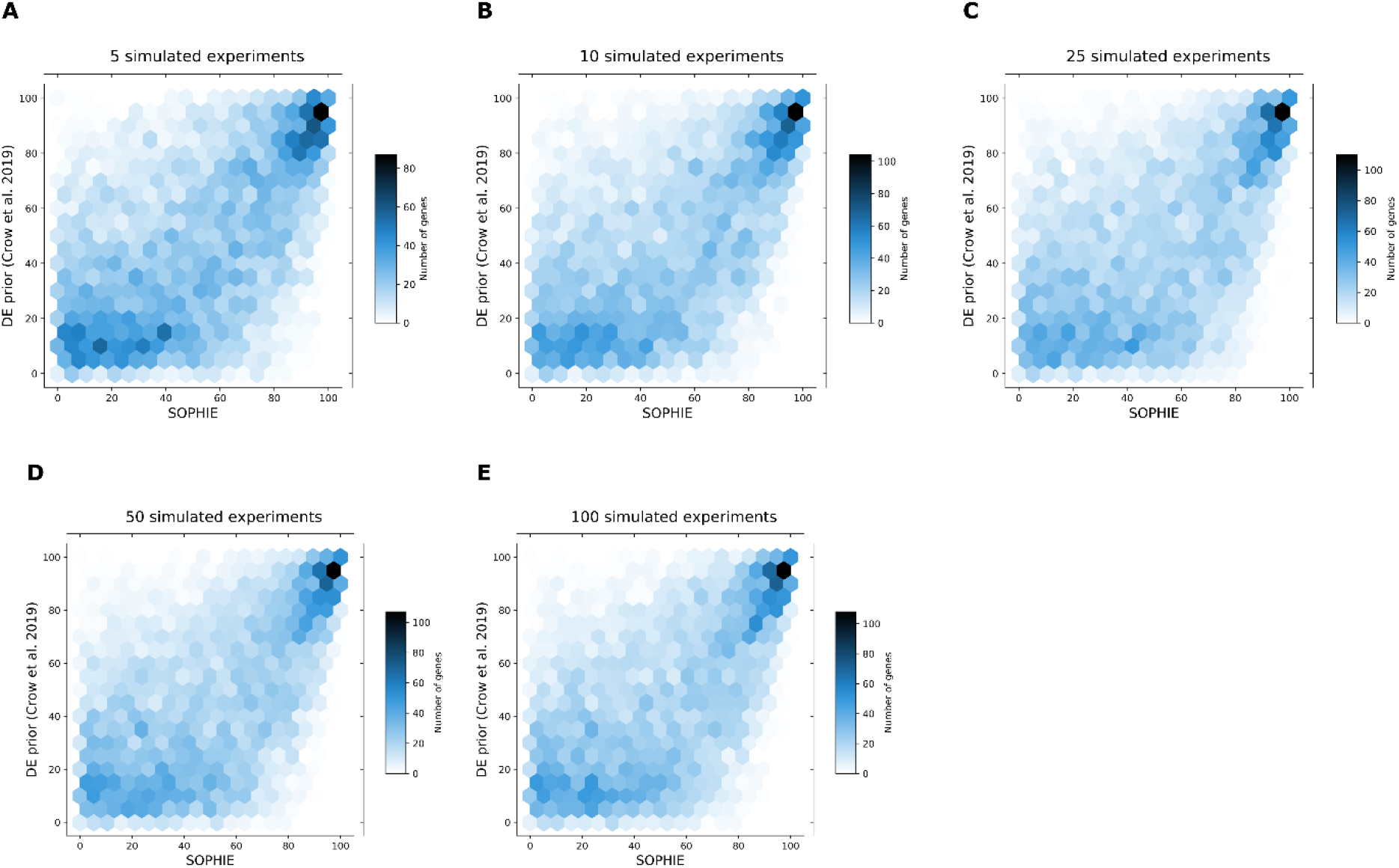
Varying the number of simulated experiments used to in the background set will yield similar results. The Spearman correlation between gene percentiles using our SOPHIE approach trained on Crow *et al.* (array) using GSE10281 as a template to generate A) 5, B) 10, C) 25, D) 50 and E) 100 simulated experiments (x-axis) versus percentiles using manually curated experiments from the same Crow *et al.* (y-axis).

**Table S1:** Human pathways associated with latent variables that contain a high (> 50%) proportion of high-weight common DEGs.

**Table S2:** *P. aeruginosa* common DEGs in that were consistent between SOPHIE trained on the *P. aeruginosa* compendium versus the GAPE curated dataset.

**Table S3:** Differential association statistics for genes regulated by the transcription factor ArgR that were found to be specific by SOPHIE.

